# Deep Learning enables reliable and comprehensive profiling of invertible promoters in microbes

**DOI:** 10.1101/2023.10.25.564076

**Authors:** Jiejie Wen, Haobo Zhang, Dongliang Chu, Xiaoke Chen, Yuxue Li, Guanxi Liu, Yuhao Zhang, Kang Ning

## Abstract

Invertible promoters (invertons) are regulatory elements found in bacteria, with inverted repeat sequences at both ends, leading to alternating changes in the expression of the regulated genes. Since invertons were present in more than 20% of bacterial genomes, while they regulated more than 5% of genes in these genomes, they are of pivotal importance for microbial functional dynamics especially when under stress. However, the prevalence of invertons, as well as the full spectrum of gene functions regulated by them, remain poorly understood. In this study, we developed DeepInverton, a deep learning model capable of accurately identifying novel inverton sequences without sequencing reads, which could profile inverton sequences from large genomic and metagenomic datasets. We conducted a pan-genomic and pan-metagenomic analysis of invertons on 68,969 bacterial genomes and 8,516 metagenome samples, resulting in a comprehensive overview of more than 200,000 nonredundant invertons and their regulated gene functional patterns. This result suggests that invertons, as a key player for bacterial adaptation to environmental stresses, are prevalent in bacterial genomes. Among the genomes analyzed, we observed a profound enrichment of invertons in pathogen such as *Bordetella pertussis*, and discovered a significant increase of inverton enrichment rates in strains associated with recent pertussis outbreaks, as well as novel evolving strains, unveiling a hidden link between the evolution of *Bordetella pertussis* and its inverton enrichment. We also utilized DeepInverton to explore inverton profiles mong human and marine metagenomes. Results revealed an unprecedented diversity of functional genes regulated by invertons, including antimicrobial resistance, biofilm formation and flagella, indicating their potential role in facilitating environmental adaptation. The *in vitro* experiments have confirmed the functions of tens of novel invertons that we have identified. Overall, we developed the DeepInverton model for exploration of invertons at unprecedented scale, which enabled our comprehensive profiling of invertons and their regulated genes. The comprehensive inverton profiles have deepen our understanding of invertons at pan-genome and metagenome scale, and could enabled a broad spectrum of inverton-related applications in microbial ecology and synthetic biology.

## Introduction

Phase variation is a well-known phenomenon in bacteria that enables frequent and reversible changes within specific transposition loci^[1]^. This mechanism maximizes the opportunities for bacteria to proactively adapt to abrupt and severe selective events, while minimizing overall genomic mutability ^[2–6]^. In bacteria, a notable process of phase variation often manifests through specific DNA regions^[9]^ that shift between two orientation, leading to altered state of gene expression. The invertible regions typically embody regulatory elements with a majority of promoters known as ‘inverton (invertible promoter)’, flanking with inverted repeat sequences ^[10]^ (**Supplementary Fig. 1**). Invertible promoters activate downstream gene expression through operator interactions in the ‘ON’ orientation. In the opposite ‘OFF’ orientation, transcription initiation is inhibited ^[10]^. Studies have reported DNA invertible promoter regions involved the promoters for numerous capsular polysaccharide biosynthesis loci ^[11–13]^, extracellular polysaccharide (EPS) ^[14]^,putative fimbriae regions ^[10, 15–17]^ and S-layer proteins and other outer surface proteins ^[5, 18–20]^. Invertons introduces phenotypic diversity into clonal populations with this reversible ON/OFF switching of individual gene expression controlled by DNA inversion of upstream promoter regions ^[21, 22]^.And the inversion events exhibit a rate that surpasses point mutations by two orders of magnitudes in the cell population ^[10, 23, 24]^. Invertons play a crucial role in endowing bacteria with adaptability to the environment, influencing essential traits related to bacterial colonization and virulence regulation ^[2–5]^.These doucumented inverton examples are associated with genes encoding pili ^[25, 26]^, flagella ^[27]^, and capsular polysaccharides ^[10, 20, 24, 28, 29]^ contribute to colonization and propagation.

Despite the documented effects of several invertons on gene expression, our knowledge of invertons in microbes still remains insufficient, especially the inverton profiles on a broad spectrum of bacterial lineages. Previous study has proposed approach known as PhaseFinder, which based on sequencing data to identify inverton^[8]^. However, due to the lack of high-throughput sequencing data spanning a large range of species, our understanding of invertons remains superficial. It is necessary to develop an all-round computational tool for effectively identifying invertons from various genomes data. Development of such a detection tool, however, is difficult due to: (1) experimental validation data of invertons is rare; (2) the sequence structure feature of inverted repetition is not sufficient to identify invertons in genomes data; (3) reliance on sequencing reads restricts the utility for tools like PhaseFinder; (4) and reads based identification approach is also slow. The read-free solution that target these bottlenecks in inverton identification might lie in AI (artificial intelligence) enabled modeling.

Here, we developed DeepInverton, a deep learning model designed for effective identification of candidate invertons independent on sequencing reads (**Figure 1**). Leveraging DeepInverton, we examined 62,291 bacterial genomes and 8,516 metagenome samples and successfully constructed a comprehensive catalog of more than 200,000 invertons. Our study unveiled an unprecedented diversity of invertons and uncovered their distribution patterns, as well as their differential function regulatory effects, across different ecological niches. Notably, pathogens such as Bordetella pertussis showed a significant concentration of invertons, which may enhance its immune evasion in host. We also confirmed function of invertons indentified by DeepInverton through in vitro experiment.. Overall, our work highlights the potential of the DeepInverton as a powerful deep learning tool for discovering candidate invertons in massive data, and forms foundation for a deeper understanding of bacterial phase inversion, as well as their applications in healthcare and environment adaptation.

**Fig. 1.**
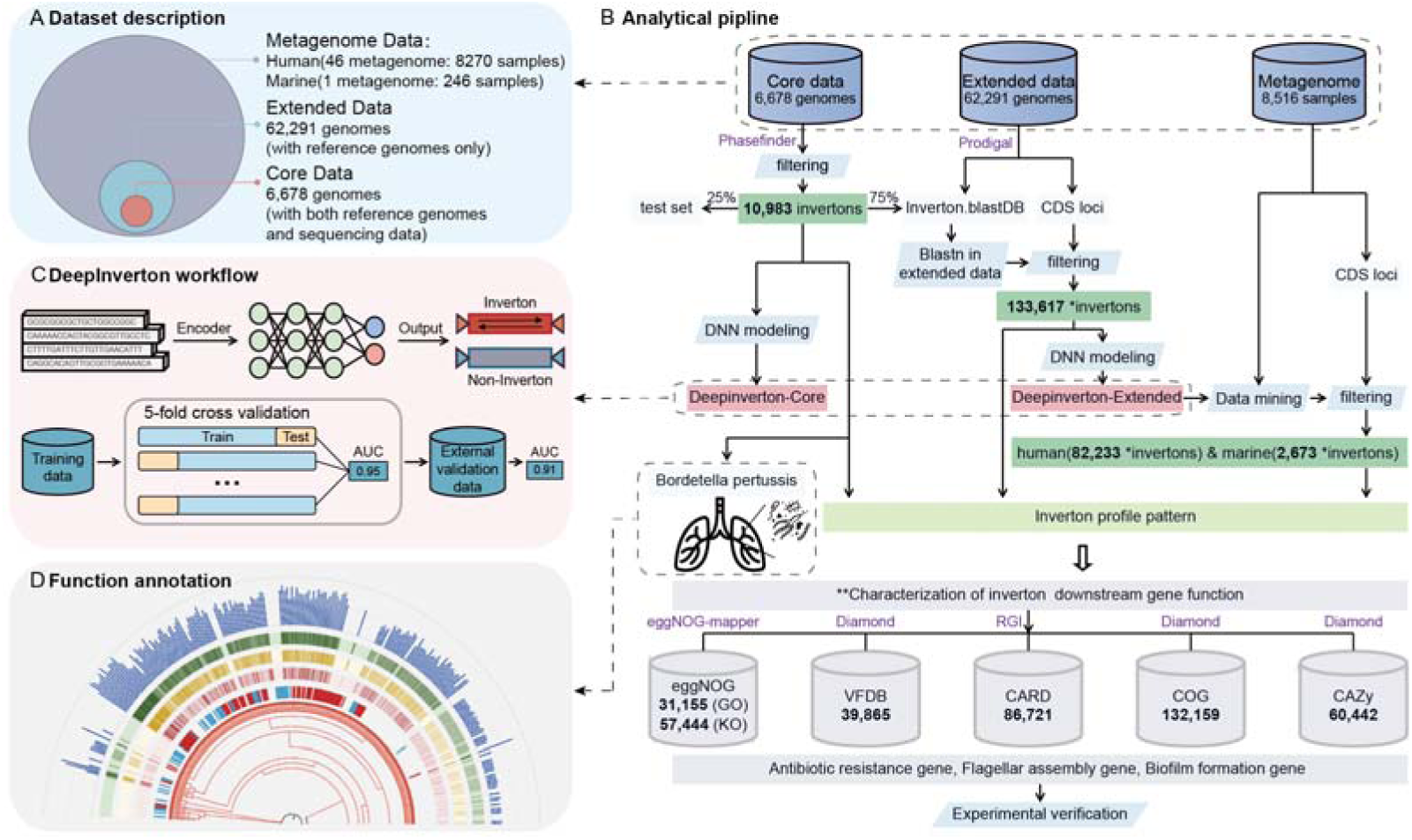
| Study workflow and dataset summary. **(A)** In this study, we started by collecting three types of datasets: core data included 6,678 available NCBI genomes from GenBank that were sequenced using Illumina paired-end sequencing, extended data included 62,291 bacterial genomes from GTDB, and metagenome data included 8270 human samples and 246 marine samples. **(B)** We mined invertons from core data using PhaseFinder and obtained 10,983 non-redundant invertons. 75% of the obtained invertons were used to searched new invertons in the extended data with BLAST. Based on these invertons, we constructed and refined neural network models for inverton identification (DeepInverton-core, DeepInverton-extended). Using DeepInverton-extended, we obtained a total of 84,906 unique invetons from metagenomic data. All the identified invertons were analyzed to characterize the inverton profile and downstream gene function pattern. *Bordetella pertussis* was investigated in great depth due to its high density of invertons. **(C)** Summary of the pipeline in building and evaluating the DeepInverton model for inverton identifying. Five-fold cross validation was applied to obtain the model performance. **(D)** A phylogenetic tree was built based on a concatenated alignment of the genomes of *Bordetella pertussis* to characterize the distribution of invertons in *Bordetella pertussis*.

## Results

### Sequence feature profiles of invertons

We compiled three datasets: core dataset, which contains 6,678 bacterial genomes with their corresponding sequencing reads curated from NCBI (details refer to **Materials and Methods**); extended dataset, which contains 62,291 non-redundant bacterial genomes (no overlap with core dataset) from the Genomic Taxonomy Database (GTDB) database^[30]^; and metagenome dataset that contains 8,516 metagenome samples from human and marine environments (Figure 1 A). The three datasets have covered a wide range of bacteria species, including 3,762 out of 4,305 known families, 169 out of 181 known phyla and 15,478 out of 19,153 known genus in the kingdom of bacteria (according to GTDB database ^[30]^, accessed as of r214), and were thus representative of most bacteria species as we currently know about.

We employed the PhaseFinder algorithm to computationally identify invertons on the core dataset and found 10,983 unique putative invertons. This search uncovered invertons in 1,664 genomes, which were found in 9 of 16 bacterial phyla (**Supplementary Table 1**). Moreover, the distribution of invertons on these genomes were highly uneven. The prevalence of invertons was higher in Proteobacteria than average, with a significant enrichment (p-value = 2.16e-07) observed at the class level specifically in Betaproteobacteria (p-value = 2.17e-07).

To further characterize sequence profiles of these invertons, we examined the top 5 phyla enriched with invertons and found significant difference in GC content (**Supplementary Fig. 2a**) and length **(Supplementary Fig. 2b**) distribution among their invertons. Interestingly, the phylum with the lowest GC content, Bacteroidetes, exhibited the longest inverton sequences, while the phylum Actinobacteria displayed a contrasting trend with the highest GC content and the shortest inverton sequence. Moreover, a comparison of GC content between invertons and their chromosomal sequences showed significant differences in 3 phyla (Firmicutes (p-value = 1.67e-07), Proteobacteria (p-value < 2.2e-16), and Actinobacteria (p-value = 1.49e-05)). An in-depth analysis of Proteobacteria in four classes including Epsilonproteobacteria, Alphaproteobacteria, Gammaproteobacteria and Betaproteobacteria, revealed notable differences among their invertons’ GC content (**Supplementary Fig. 2d**) and discrepancy compared with their respective chromosomal GC content (**Supplementary Fig. 2f**). While there were no significant difference observed in terms of inverton length (**Supplementary Fig. 2e**). (**Supplementary Fig. 2e**).

To decipher distribution of orientation state among invertible promoters, we calculated the proportion of inverton sequences supporting the reverse orientation within the core dataset. We found a bimodal distribution of Pe_ratio (proportion of the reverse orientation sequences, details refer to **Supplementary Table 5**) around 0.02 and 0.5 (**Supplementary Fig. 3A**), with the peak of 0.02 showing a higher density. This result may suggests that invertons are typically silenced under regular conditions. However, the inversion events could rapidly increase to active specific gene expression in response to intense interspecies competition or environmental selection pressures, resulting a ratio approaching 50% between inversion and non-inversion. Furthermore, we found that the invertible promoter motif patterns were consistent with previously reported study^[8]^, supporting the high reliability of inverton sequences provided by the core dataset (**Supplementary Fig. 3**).

### Profiling of invertons across a phylogenetically broad spectrum of bacteria using 62,291 genomes

To uncover the comprehensive distribution of invertons in bacteria, we searched the extended dataset, which covered all annotated genus in the GTDB and could represent one of the most wide range of known bacteria (details refer to **Supplementary Table 5**). Based on BLAST sequence search algorithm (details refer to **Supplementary Table 5**), 133,617 putative invertons were obtained (**Materials and Methods, Supplementary Table 2**),,which showed a consistence in consensus motif with the invertons identified from the core dataset (**Supplementary Fig. 3D**)

Next, we constructed a phylogenetic tree with explored species (involved in both core dataset and extended dataset) (**Figure 2A**) based on their genomes. Phylogenetic analysis of this tree revealed that the invertons are extensively distributed, but they were predominantly concentrated within 4 phyla (Proteobacteria, Actinobacteria, Firmicute, and Bacteroidete) as well as 5 genus (*Pseudomonas*, *Bordetella*, *Bacteroidetes*, *Mycobacterium*, and *Staphylococcus*) **(Figure 2A, labels 1-5**). We readily detected that the species in extended dataset demonstrated a homogeneous pattern with the core dataset, verifying the scalability of mining the extended dataset according to the core dataset. Additionally, invertons are observed to be prevalent in consistent species in the two datasets. We have also discovered novel invertons on several lineages which were exclusively present in extended dataset (**Figure 2A, labels 6-7**), such as Desulfobacterales, Campylobacterales and Myxococcota **(Figure 2B**), supporting the “more genomes, more invertons” hypothesis on global scale(**Supplementary Figure. 4**). On the other hand, for certain species such as Bordetella pertussis that contain hundreds of strain genomes, the saturation curve has almost reached the plateau. Interestingly, among the top 10 species with the highest IPK (Number of Invertons Per Kilobase of genomes), 8 of them correspond to pathogenic bacteria. As the mean IPK value of the topN species decreased, there was a corresponding downward trend observed in the proportion of pathogens in these species (**Supplementary Fig. 5**). This enrichment of invertons in pathogenic species, providing an opportunity to gain new insights into the evolution of host-pathogen-inverton interactions.

**Fig. 2.**
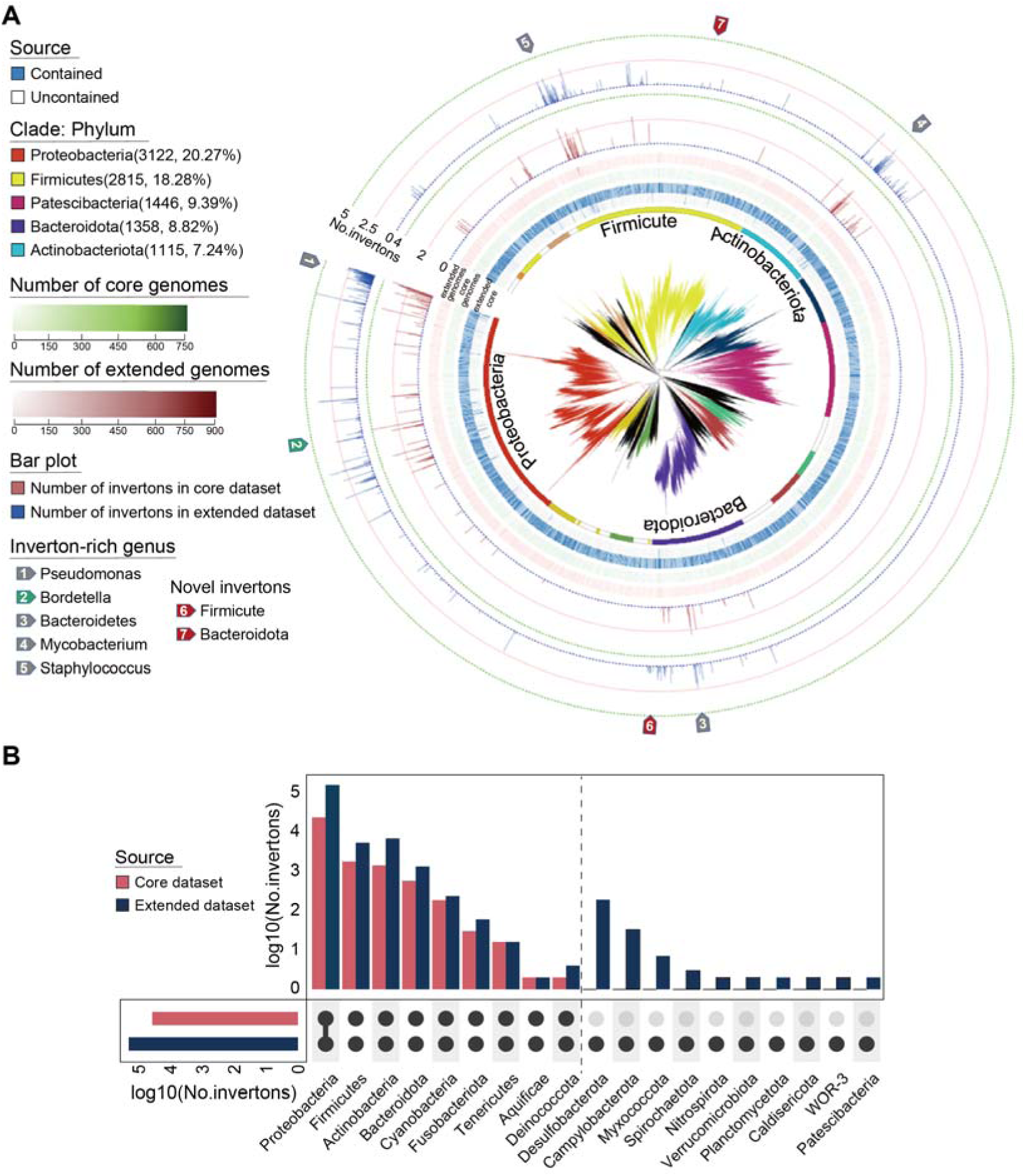
| The global distribution pattern of identified invertons. **(A)** A phylogenetic tree was built based on a concatenated alignment of the species genomes in core and extended data. Label displays the family annotation and different colors indicate the top 12 phyla level species (top5 phyla were labeled). The inner two strip charts indicate the data source of the species (core data or extended data). The next two strip charts indicate the number of species genomes in core data and extended data. The next two bar plots indicate the number of invertons in core dataset and extended dataset. The area indicated by label (1-5) shows the enrichment of the number of invertons in certain related species at the same phylum level, and label (6-7) indicates that the invertons were identified only in the extended data and not found in the core data. **(B)** Number of invertons in different bacteria phyla from core dataset and extended dataset. Invertons have been found in the species that do not appear in the core dataset (right side of dotted line), such as Desulfobacterota, Campylobacterota, Myxococcota, Spirochaeta, etc.

### Functional analysis of downstream genes regulated by invertons

To provide a systems-level functional regulation snapshot of invetons, functional annotation was created on the genes regulated by invertons from both core dataset and extended dataset. Firstly, on the core dataset, the annotation included 11,033 GO annotations and analysis of conserved genes showed that were primarily associated with stress adaptation, environmental interaction and membrane formation (**Supplementary Fig. 6A**). KEGG pathway analysis revealed function enrichment related to two-component system, membrane synthesis, flagellar assembly, and drug resistance (**Supplementary Fig. 6B**). Additionally, the annotation results in virulence factor database (VFDB) and the Comprehensive Antibiotic Resistance Database (CARD) (**Supplementary Table 3**)showed abundant functions associated with bacterial motility and antibiotic resistance, supporting the GO and KEGG findings. Consistent with core dataset, the functional genes regulated by invertons are primarily associated with membrane synthesis and stress resistance in extended dataset (**Supplementary Fig. 7**). Overall, these results hint potential relationships between invertons and host interaction with the external environment, as well as their role in host’s stress resistance.

### Characterization of the Inverton profile in *Bordetella pertussis*

A total of 6,350 invertons were identified in *Bordetella pertussis* (the pathogen of whooping cough) from core dataset, and with an average of 15 invertons per strain among the 426 strains under this species. Functional annotation of genes regulated by these invertons revealed that 4,325 regulated antibiotic resistance genes on average, accounting for more than 50% of all inverton regulated genes.

The majority of invertons are We constructed a phylogenetic tree with the 426 genomes of *B. pertussis* from the core dataset, in addition to 127 genomes of various closely related species spanning different taxa levels (**Figure 3a**). Results on this phylogenetic tree has confirmed again that *B. pertussis* exhibited a higher abundance of invertons (p-value = 1.38e-24) compared to other species. Moreover, we compared the inverton profiles of *B. pertussis* strains categorized by whooping cough outbreaks (before and after 2011 ^[31, 32]^, see **Materials and Methods** for details). It was found that the number of invertons in the *B. pertussis* strains from the outbreak after 2011 was significantly higher than those from the outbreak before 2011 (p-value = 0.0041) (**Figure 3b**).But there was no significant difference in the number of invertons regulated functional genes categorized as motility genes like flagella and pilus synthesis (**Figure 3c**) and antibiotic resistance genes (ARGs) (**Figure 3d**) between the two groups. These increased invertons might assist *B. pertussis* strains to escape existing vaccines or therapeutic drugs, possibly inducing to new outbreaks of pertussis.

**Fig. 3.**
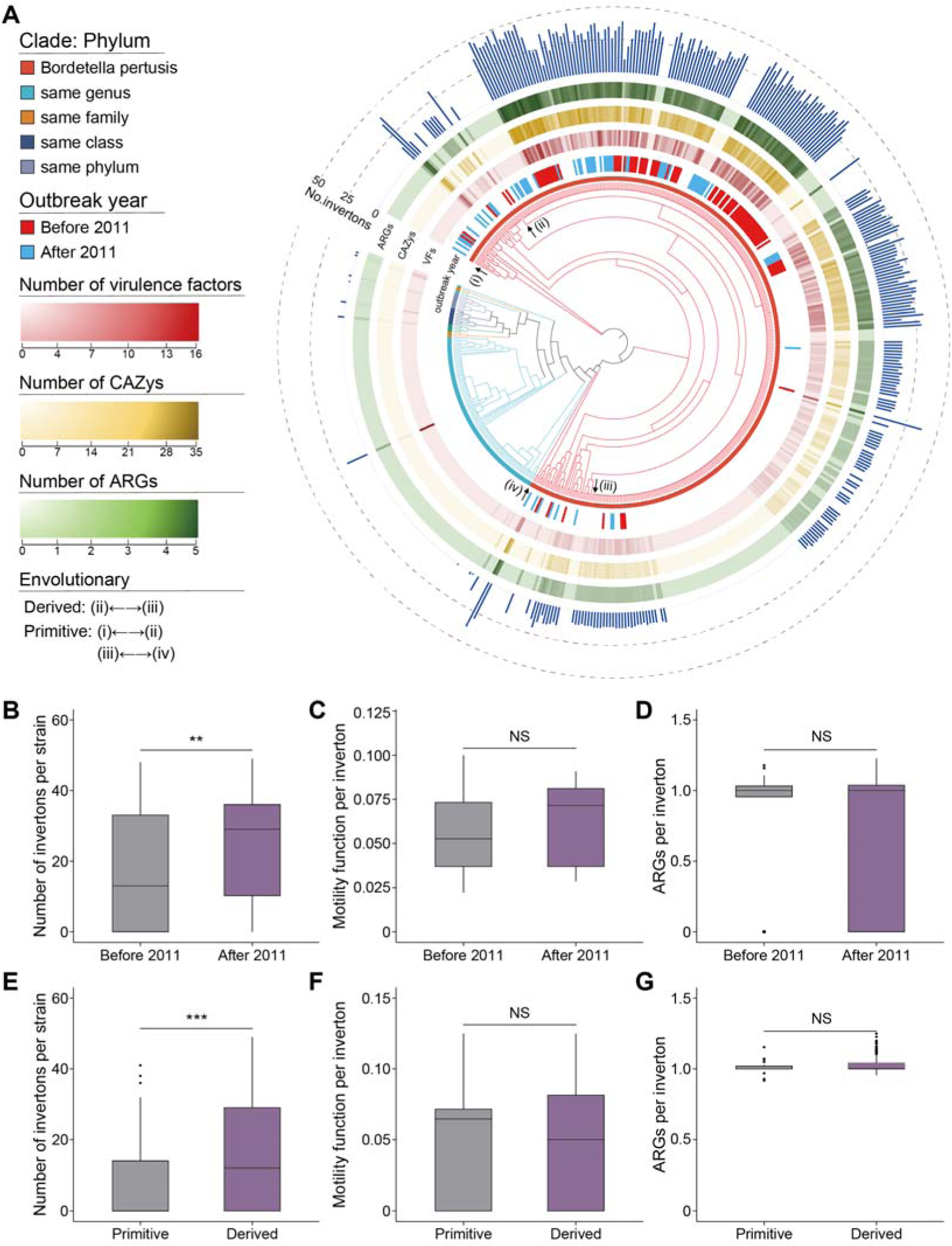
| Characterization of inverton profile in the *Bordetella pertussis*. **(A)** A comprehensive phylogenetic tree was constructed by integrating the genomes of all *Bordetella pertussis* strains in the core dataset, alongside genomes from taxonomically related species within the same genus, family, order, class, and phylum. The depicted phylogenetic tree provides a hierarchical representation of evolutionary relationship. The color gradient, ranging from red to blue, demarcates distinct temporal cohorts, with red denoting strains originating “after 2011” and blue indicating those predating this period. The next three strip charts indicate the cumulative count of inversion-regulated virulence factor genes, carbohydrate-active enzyme genes, and antibiotic resistance genes. The outermost blue bar graph elucidates the distribution of inversions across the genomes. Taxonomic classification of *Bordetella pertussis* strains is determined based on evolutionary lineages, specifically delineating derived strains (denoted by labels (ii) to (iii)) and primitive strains (denoted by labels (i) to (ii) and (iii) to (iv)). **(B)** The comparison in distribution of invertons and its regulated function among different groups categorized as “before 2011” and “after 2011,” as well as “primitive” and “derived”.

Furthermore, we found similar characteristics in the evolution of *B. pertussis* strains. Compared to primitive strains (**Figure 3a**), the more evolutionarily derived strains showed a higher number of invertons (p-value = 1.709e-07) (**Figure 3e)**. These invertons regulated plentiful functional genes related to bacterial motility (**Figure 3f**) and ARGs (**Figure 3g**), which potentially enhanced the survival rate in extremely harsh environments. For instance, the emergence of the evolutionarily derived strain NZ61 broke out in 2017 ^[32]^, an inverton located at position 1,048,412 to 1,048,505 was identified **(Figure 4a**), which regulated the synthesis of the flagellar transcriptional regulator FlhD. This regulatory mechanism had an impact on the expression of multiple genes involved in flagellar synthesis, including the flagellar motor protein MotA located downstream. MotA serves a crucial function in the assembly and functioning of the flagellar motor, facilitating the energy-driven rotation of the flagella. Furthermore, on the strain I518 broke out in 2012 ^[31]^, we identified a inverton at position 2,070,216 to 2,070,429 that regulate the *sul gene* **(Figure 4a)**, which encodes a sulfanilamide-resistant enzyme. This enzyme could improve the ability to metabolize sulfanilamide drugs or modulate their cellular uptake and efflux mechanisms in bacteria. Overall, the increasing frequency of invertons in BP might help the host to better adapt to environment, for example the enhancement of the immune evasion to antibiotic treatment.

**Fig. 4.**
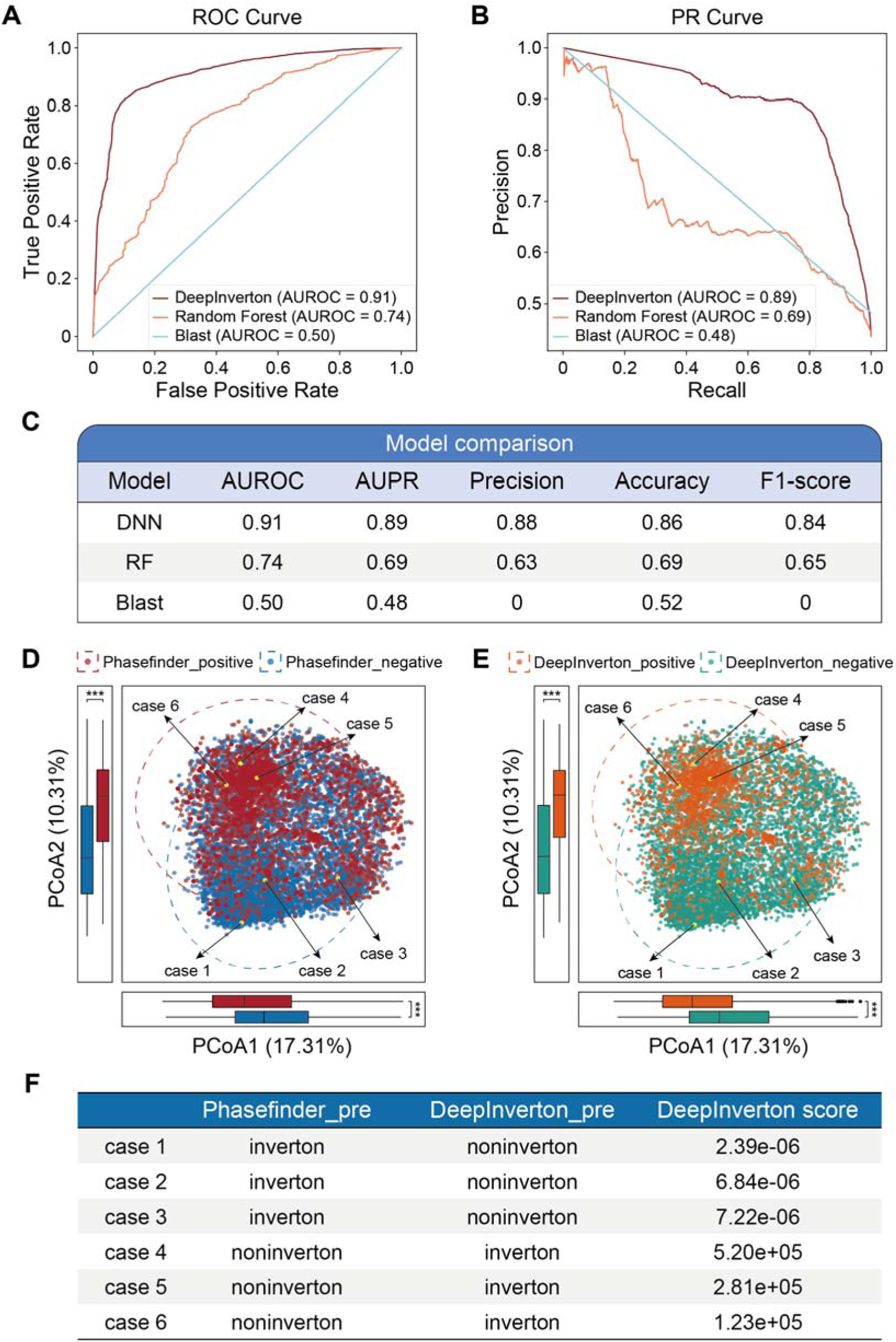
| Comparisons of DeepInverton with other methods in identifying invertons on the external validation dataset. (A, B) Area under the receiver operating characteristic curves (A) and area under the precision-recall curves (**B**) show the performance comparison of DeepInverton (red), RF (orange), and BLAST models (blue). **(C)** Performance metric details of the trained models for identifying invertons from non-invertons using sequence data in the external validation set. (**D, E**) External validation set sequence 3-mer PCoA analysis. PhaseFinder positive (red) and PhaseFinder negative (blue) are clustered into two categories, in which Phasefinder positive set contains a set of sequences identified as invertons by Phasefinder, while Phasefinder negative set contains a set of sequences identified as non-invertons by Phasefinder. The same definitions apply for DeepInverton positive (pink) and DeepInverton negative (green) set. The “inverton positive cluster” refers to set identified by either Phasefinder or DeepInverton, while the same definition apply for the “inverton negative cluster” (refers to **Supplemetnary Table XXX** for details). (**F**) Invertons prediction for case 1-6 by PhaseFinder and DeepInverton.

### DeepInverton enabled candidate invertons identification at large scale without sequencing reads

We developed DeepInverton, a sequence based deep learning-enabled method, for inverton identification. The development of this predictive tool was motivated by the hypothesis that sequencing-data-independent inverton prediction could facilitate biologically inspired prospection for candidate invertons bacteria on broader scale (**Figure 1c**). The DeepInverton models took as input a fragment of sequence, and could output an inverton candidate that is composed of three different parts: the inverton itself and its flanking reverted repeat sequences. Using the curated training set derived from core or extended inverton datasets, we obtained two models known as DeepInverton-Core and DeepInverton-Extended, respectively. The performance of DeepInverton models has been evaluated on an external test set, which was pre-allocated from the core inverton dataset and entirely independent of the model construction (**Figure 1b**).

We found that both DeepInverton-Core and DeepInverton-Extended exhibited superior classification performance, achieving an AUROC (Area Under the Receiver Operating Characteristic curve) over 0.90. Compared to DeepInverton-Core, DeepInverton-Extended exhibited a potential of better adaptability and robustness in identifying invertible regions from diverse species due to the broader species coverage and inclusion of more inverton sequences. Consequently, DeepInverton-Extended was selected as the default model for the identification of invertons in metagenome dataset.

A comparison with other methods using the same external test set highlighted the superiority of DeepInverton as regard to prediction accuracy (**Figure 4**). In contrast to DeepInverton, the Random Forest model demonstrated poorer performance, achieving an accuracy of 0.69, an AUROC of 0.74 and an AUPRC (Area Under the Precision-Recall curve) of 0.68. For the traditional BLAST methodology, its contribution to the identification of invertons was found to be negligible, with an AUROC and AUPRC both less than 0.5. Notwithstanding the high sensitivity of BLAST, reaching to XXX, DeepInverton, while maintaining a comparable sensitivity, excels in the expeditious and robust identification of a substantial quantity of reliable invertons.

Overall, these results support that DeepInverton is an effective and robust approach for Inverton identification from sequence data.

Interestingly, numerous promoter sequences featured with inverted repeats could not be identified by the DeepInverton. To explore the dissimilarities between inverted repeats promoters^[35]^ (IRP, regular promoters with inverted repeat sequences) and invertible promoters, we examined putative promoter sequences flanked by inverted repeats from the genomes within the core dataset. Some of these promoters were further identified as invertons by PhaseFinder and DeepInverton. As a result, our analysis revealed that considerable differences in the 3-mer features of IRP when compared to the predicted invertons by PhaseFinder and DeepInverton (**Supplementary Fig 8**).These results supported notion that PhaseFinder and DeepInverton utilize characteristics beyond the mere sequential structural feature of inverted repetition for identifying invertons from the massive and intricate genome sequences.

In contrast to the PhaseFinder program, we have emphasized the superiority of DeepInverton in design. First of all, DeepInverton solely relies on the original inverton sequence and its flanking inverted repeat nucleotide sequences as the input information, enabling inverton discovery for almost all bacterial species once their genomes or partial genomes are available. It is also noteworthy that DeepInverton effectively circumvent the computationally intensive steps needed by PhaseFinder, which include mimics involving diverse inverton orientations within a species’ genome or comparisons with the metagenomic data ^[8]^. Most importantly, DeepInverton has demonstrated a comparable performance while only utilizing the inverton sequence features (without sequencing reads). We have discovered that DeepInverton identified 80.18% of the invertons previously determined by PhaseFinder in the external test set (**Supplementary Fig. 9a**). DeepInverton showed the high fidelity with a low “false positive rate” less than 10% (**Supplementary Fig. 9a**).

Additionally, our analysis revealed noteworthy disparities in 3-mer nucleotide infromation between inverton and non-inverton, as determined by either PhaseFinder or DeepInverton (**Figure 4D, E**). This observation underscored that DeepInverton possessed a comparable capacity to PhaseFinder in accurately segregating two distinct sets of sequence features for inverton and non-inverton. Moreover, we observed that three sequences (**Figure 4F, case 1,2,3**), which were classified as non-invertons by PhaseFinder but identified as invertons with high probabilities by DeepInverton, exhibited similar 3-mer feature profiles to the center of the inverton positive cluster determined by either PhaseFinder or DeepInverton (**Figure 4D**). Conversely, another three sequences (**Figure 4F, case 4,5,6**), which displayed similar 3-mer feature profiles to the center of the inverton negative cluster determined by either PhaseFinder or DeepInverton (**Figure 4E**), were identified as non-invertons by DeepInverton but opposite by PhaseFinder. Thus, DeepInverton These findings suggest that many of the invertons identified by DeepInverton but missed by PhaseFinder were likely to be genuine invertons.

### Identification of 84,906 candidate invertons from 8,516 metagenome samples

To understand the enrichment patterns of invertons across diverse ecological niches, we leveraged DeepInverton to explore the invertons on the metagenome dataset collected from human and marine microbiome data ^[36, 37]^. The human microbiome data consisted of 46 studies (**Supplementary Table 4**) encompassing 8,270 human samples spanning diverse populations, different host ages and multiple body sites. We also incorporated a marine microbiome data including 246 marine samples, which provides a broad and diverse spectrum of genomic background for inverton analysis. A stringent criteria was executed to ensure that the identified invertons display a sufficient probability of being true (**Materials and Methods**), and a total of 84,906 unique putative invertons were ultimately obtained. Among these, 82,233 putative invertons were derived from the human microbiome, from which we found that genes regulated by invertons are assocaited with signal transduction and biofilm synthesis (**Supplementary Fig. 10**). For marine microbiome, we have identified 2,673 candidate invertons. Interestingly, despite of functional enrichment in extracellular region, signal transduction and biosynthesis of biofilm, we have also found that a prevalence of inverton downstream genes related to the flagellum, indicating another strategy against environmental stress owned by bacteria in the marine ecosystem.

We systematically examined the distribution of invertons across the four datasets including core dataset, extended dataset, human metagenome dataset, and marine metagenome dataset on the phylogenetic tree (**Figure 5**). We observed a consistent and statistically significant predominance of invertons within the phylum Proteobacteria (p-value = 2.12e-09). There was a unique distribution pattern of inverton emerged in the human microbiome, demonstrating a remarkably extensive presence of invertons in Firmicutes and Bacteroidetes, even higher than the Proteobacteria phylum (p-value = 2.12e-09). Furthertmore, invertons were notably prominent in pathogenic bacteria, particularly within the genus Neisseria (p-value = 4.22e-04). This observation highlighted the potential role of invertons in the genomic repertoire of pathogenic bacteria.

**Fig. 5.**
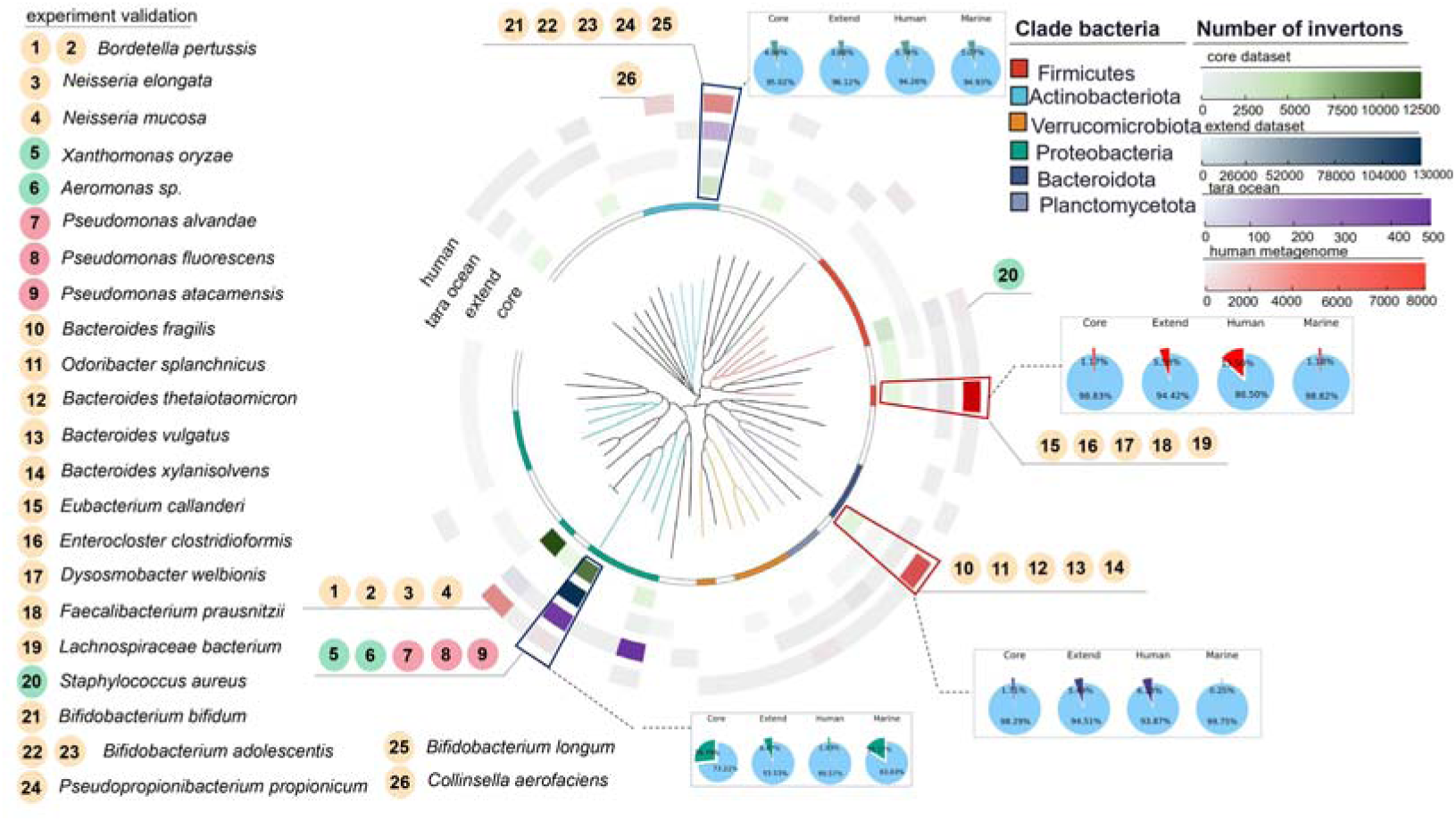
| Distribution of the count of the invertible regions at different bacterial class levels across the four datasets. Evolutionary trees were constructed based on representative genome sequences from each class. The heatmap, from inner to outer layers, displays the distribution of invertible region counts for the “core dataset”, “extended dataset”, “human metagenomes”, and “marine metagenomes”. We performed external experimental validation on the labeled sequences from 1 to 26. Among them, the green labels (5, 6, and 20) correspond to case-1, case-2, and case-3, while the red labels (7, 8, and 9) correspond to case-4, case-5, and case-6. The other sequences (labeled with orange color) were identified as invertons by DeepInverton from the metagenomic data with high fidelity.

Next, we confirmed that although the DeepInverton’s prediction results were influenced by the training set, it had fairly good extendibility for inverton identification. To evaluate the differences between identified invertons from the two metagenome datasets and core dataset, we performed sequence alignment using BLASTn ^[33]^. This analysis uncovered that a subset of 12,077 putative inverton sequences identified through the DeepInverton exhibited notable unsimilarity (p-value < 1e-05) to the invertons present in the core dataset. And the distinctive pattern in the human microbiome can be attributed to the predominant presence of invertons in Proteobacteria (65.41%), Firmicutes (16.88%), Actinobacteria (8.48%) and Bacteroidetes (7.06%), which are known to constitute the most prevalent bacterial populations in the human intestinal ecosystem. However, the DeepInverton was trained with the invertons from the core dataset and extended dataset, both of which have lower enrichment of invertons within Firmicutes (6.65%), Actinobacteria (5.24%) and Bacteroidetes (2.12%). Nevertheless, DeepInverton has identified plenty of candidate invertons within these bacterial species (32.42%) in the human microbiome samples. These findings substantiated the exceptional ability of DeepInverton to discover novel candidate invertons beyond the known repertoire.

Collectively, by leveraging the deep learning architecture, DeepInverton exhibited a remarkable capacity to uncover previously unexplored invertons from diverse genomes. It could broaden our understanding of genomic landscapes and expand the scope of inverton research. To enable its convenient use by the scientific community, the more than 200,000 candidate invertons resource — which include detailed genomic information of core invertons, extended invertons and metagenome invertons — has been made openly accessible alongside taxonomic and functional annotation at the Inverton Database http://inverton-db.aimicrobiome.cn/. The DeepInverton model has also been made publicly accessible at https://github.com/HUST-NingKang-Lab/DeepInverton.

### Profound enrichment of inverton after the Cambrian explosion event

An unbalance were observed in the distribution of inverton in bacteria based on the phylogenomic analysis of invertons (**Figure 2A**, **Figure 6**). To understand this phenomenon in light of the evolutionary relationships, we focused on the evolutionary dynamics of invertons during billions of year on genome-wide scale. The divergence times were obtained from the Timetree distributed across 2458 genera fully representing 99.64% of inverton amount in bacteria. A growth of invertons observed after ∼500 million years ago (Ma), coinciding with the ‘Cambrian Explosion’ event occurred approximately from 540 to 520 million years ago with the rapid increase in species diversity and abundance.

**Fig. 6.**
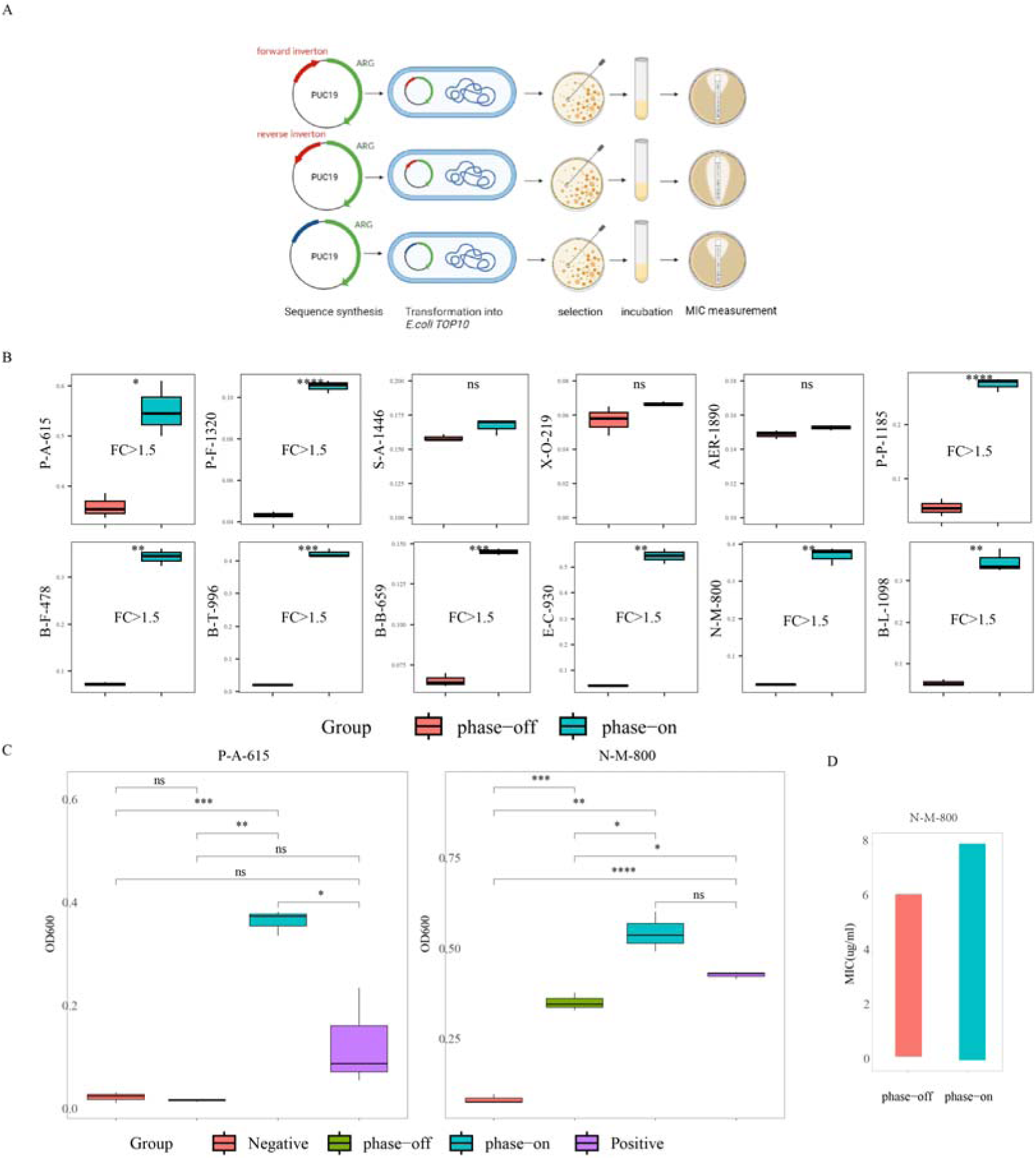

Invertons have extended their host lineages from 12.39% to 20.71%, and the statistical analysis revealed a significant increase in the average number of invertons per genera after this time point (**P-value = 0.016, Supplementary** Fig.11 **A**). Additionally, the diversity of genes regulated by invertons showed the types of genes rising from 2434 to 4018 following the Cambrian (**Supplementary Fig. 11B**). Further in comparison to the function of these genes, we found an enrichment of after-Cambrian genes associated with response to stimulus and membrane (**Supplementary** Fig.11 **C**) which are crucial for the bacterial survival, reproduction, immune defense, and evolutionary genetics. The possible reasons for these growth phenomena are bacteria require the adaptive strategies to strengthen their adaptability to abrupt environment and outperform other bacterial species in the competition for nutrition after the Cambrian with more diverse taxa occupied distinct ecological niches and resources.

### Experimental validation of candidate novel invertons identified by DeepInverton

We then synthesized the representative candidate novel invertons identified by DeepInverton for verification and characterization. Both ON and OFF orientation sequences were synthesized to represent the regulatory forms of the invertons in vivo. Two candidate invertons were discarded due to the failure of syntheses or experimental manipulation. We examined the remaining 21 (including the case 4-6, **Figure 4F**) candidates’ regulation of downstream antibiotic resistance genes (ARGs) at the antibiotic exposure. This verified 9 candidates with inverton’s regulatory function under a stringent threshold (that is, the survival of bacteria with ON or OFF orientation sequences showed a significant difference with P-value less than 0.05 and Fold-change higher than 1.5), representing a positive rate of 42.9% (9/21). Notably, case4 and case5 are among the 9 verified candidates, supporting the prediction by DeepInverton. For case6, though it was not among the 9 verified candidates, it met the P-value threshold, and was highly likely to be a true positive. On the other hand, a negative set of three candidate non-invertons (case 1-3) were examined, of which all three demonstrated no significant difference in the bacteria survival under ON or OFF orientation sequences. These results confirmed DeepInverton’s prediction, suggesting the high reliability of our method in inverton identification.

We further examined the bacteria survival under known ARGs active (positive control) or silenced (negative control) to explore the invertons’ regulation of gene expression. Specifically, we found a significantly improved survival of bacteria exposed to penicillin under the expression of streptothricin acetyltransferase (*SatA*) gene, which has been previously reported to be resistant to penicillin. Bacteria with ON orientation sequences showed similar survival condition to positive control (OD_600_ values around 0.01), and the OD_600_ value for OFF orientation sequences group was close to those of the negative control (OD_600_=0.4 and 0.1 for OFF orientation sequences and negative control, respectively). For another ARG against erythromycin, *macB* gene presented in *Neisseria mucosa*, a pathogen that could cause endocarditis. It demonstrated a consistent expression pattern regulated by inverton sequence orientation. Additionally, the result of MIC determination also supported the above findings, exemplified on *macB* gene (Figxx). These results establish that the sequence orientation change effectively regulates the expression of ARGs and could potentially affect the escape of some pathogens to antibiotic therapy.

## Discussion

Invertible promoters are pivotal elements that reversibly regualte gene expression, but the global profiles of bacterial invertible promoters were still untapped. Here, we developed the deep learning model, DeepInveton, to effectively identify putative invertons without the dependent on sequencing reads, enabling exploration on large-scale resources. In an unprecedented effort to unearth more than 200,000 putativate invertons on extensive genome and metagenome microbime communities, we have uncovered the comprehensive catalog of invertons spanning the sequence profiles, distribution pattern and function annotation of the inverton downstream genes.

DeepInverton surpasses exsiting alternative approaches in precisely and efficiently detecting invertons without relying on sequencing data. Despite the model’s training data contained only a specific set of bacteria, it had exhibited useful applications in identifying candidate invertons across diverse species and niches. Furthermore, DeepInverton performed the superiority over PhaseFinder, it could soley pick up certain potential invertons that PhaseFinder had missed. The follow-up experimental investigations demonstrated that our method can discover diverse and novel invertons. DeepInverton should be valuable to the research community, as it promoted the model-oriented experimental validation for inverton knowledge discovery. DeepInverton has facilitated the accurate and rapid dicovery of inverton on a large scale, and it may motivates researchers to assess a list of candidate invertons from massive data, providing targets for further exploration.

Our systematic exploration of inverton distribution and genomic correlates revealed that potential invertons were prevalent in pathogenic bacteria including *Bordetella pertussis*, *Neisseria sicca*, *Neisseria mucosa* and *Neisseria meningitidis*. These invertons were found to regulate genes primarily associated with bacterial motility, biofilm assembly and antibiotic resistance, suggesting the potential of invertons in the pathogen’s evolutionary strategies in medical intervention environmental adaptation. Such knowledge and discoveries also provide a fresh perspective on our understanding of bacterial adaptation, host-pathogen interactions and the progression of infectious diseases.

The predicted functions of genes regulated by inverton likely reflect the specific environmental adaptation strategies.Our results revealed the enrichment of antibiotic resistance, biofilm formation and flagella, while the possible machenisms remain illusive..The maintenance of inverton regulatory mechanisms may serve as a bet-hedging strategy ^[43]^, enabling bacteria to rapidly switch between different phenotypic states. Conversely, continous expression strategy evolves more slowly because it requires the ongoing energy consumption, which is not conducive to bacterial survival in energy-deprived harsh environments. Thus, inverton that is a better option for bacteria to flexibly regulate the gene expression can confer a selective advantage in diverse environments, host defenses, nutrient availability, and competition with other microbes. Although this hypothesis is difficult to test experimentally, our results are in line with the computationally predicted on large-scale genomic and observed of inverton content on evolutionary timescales and variation with survival resource availability.

Despite this work has uncovered certain mysteries of inverton, this is just the tip of the iceberg for the complete comprehension of this strategy. In the current study, we did not incorporate longer inverton sequences into DeepInverton, even though less than 300 bases represent the typical length distribution of invertons in majority of bacteria (**Supplementary Fig. 9b**). More importantly, it is always imperative for researchers to dedicate their efforts to unmask inversion strategy, comprehend the inverton evolution process and apply this mechanism to assist healthcare.

In conclusion, to our knowledge, this work offers the largest inverton resource and provides comprehensive insights into the distribution and functional regulation of invertons across a broad spectrum of bacteria inhabiting host-associated and environmental niches. This study developed the high-fidelity utility of integrating DeepInverton with massive microbiome data for inverton exploration, prepresenting a ‘re-purposing’ application for the extensive researches in environmental and medical metagenomics with large-scale datasets to discover a fraction of the functional ‘dark matter’. The DeepInverton model, together with more than 200,000 unique candidate invertons that we have identified, could server as a foundation for deeper understanding of invertons, and future exploration about bacterial adaptation to stressful environment. They could also provide a resourceful toolkit with applications spanning healthcare, environment and industry, such as providing possible module for synthetic biology to allow more flexibility in the bacterial genetic circuit.

### Key points

(1) A deep learning model DeepInverton has been built for mining of candidate invertons without the need of sequencing reads, and with high fidelity.
(2) A catalog of sub-million canditate bacterial invertons and their regulated genes has been established, while tens of novel invertons were experimentally validated, suggesting that invertons are prevalent in bacterial genomes.
(3) Pathogenic bacteria exhibited enrichment of invertons, exemplified by *Bordetella pertussis*, which has a profoundly enrichment of invertons, probably leading to their high adaptability to stresses.
(4) Mining of human and marine metagenome has revealed an unprecedented diversity of functional genes regulated by invertons, including ARGs, flagellar and biofilm assembly, most likely be responsible for environmental adaptation.

## Materials and methods

### Datasets involved in this study

Firstly, we downloaded 6,678 available genomes from Genebank that were sequenced using Illumina paired-end sequencing with data deposited in the NCBI Short Read Archive as the core dataset. PhaseFinder (v1.0)^[8]^ was employed to identify putative invertible regions in the genomic assemblies with the default parameters. We filtered the results by removing the invertible DNA regions with less than 2 reads supporting the R orientation from the paired-end method, or a Pe_ratio below 1%. Furthermore, the invertible DNA regions containing or overlapping coding sequences (CDS) were eliminated. A total of 10,983 inverton sequences are obtained in the core dataset.

Secondly, we also included an extended dataset comprising of 62,291 bacterial genomes obtained from the genome taxonomy database (GTDB, release207)^[30]^ to further supplement the core dataset. We used 75% sequences of the core dataset to build a homology alignment database, and then searched the nucleotide sequences of these 62,291 genomes using BLASTn (v2.13.0)^[33]^ with the following parameters: perc_identity=90, evalue=1e-5. We also removed the sequences aligned to less than 95% of the length of the target inverton sequence, and removing the sequences contained in core data. Finally, 133,617 sequences were acquired in the extended dataset.

For the metagenome dataset, 46 metagenome dataset that contains 8,516 metagenome samples from human and marine environments (Figure 1 A).

### Gene function annotation of sequences regulated by the invertons

We extracted nucleotide sequences with a length of 2000 base pairs downstream of the invertons for gene identification and functional annotation. The extracted nucleotide sequences were annotated to the KEGG and GO databases using the eggNOG-mapper (v2.1.6)^[44]^ tool with default parameters. For Gene Ontologies (GOs), we first obtained microbiology-related GO-slims from goslim_metagenomics.obo (accessed on November 11, 2022) provided by geneontology.org. Then, we mapped the gene orthologs (GOs) generated by eggNOG to different subgroups of Molecular Function (MF), Biological Process (BP), and Cellular Component (CC). Finally, the counts of GO terms in the three GO categories were calculated, and the top five GO terms in each GO domain were obtained. For KEGG metabolic pathways, we mapped the KEGG orthologs obtained from the annotation results of eggNOG to the KEGG modules providing by the KEGG BRITE database (accessed on May 5, 2022). The abundance of each KEGG Orthology (KO) was calculated by summing the abundance of genes annotated to that particular KO, and the top 20 KEGG maps were obtained.

Furthermore, we have also annotated the inverton downstream gene function to other databases, including Comprehensive Antibiotic Resistance Database (CARD, 3.2.6 January 2023 release)^[45]^, Virulence Factor Database (VFDB, 2022)^[46]^, Cluster of Orthologous Groups (COG, 2020) and Carbohydrate-Active enZYmes Database (CAZy, 2020)^[47]^. Antibiotic resistance genes (ARGs) were annotated by querying the de-replicated gene sequences against the CARD using Resistance Gene Identifier (RGI, version 3.2.6) with the loose model (–include_loose). Virulence factors were annotated by aligning gene sequences against the VFDB with DIAMOND blastx (e-value threshold of 1×10−5, v2.1.1.155)^[48]^. The known protein families were annotated by aligning gene sequences against the COG with DIAMOND blastx (e-value threshold of 1×10−5). And the carbohydrate-active enzymes were annotated by aligning gene sequences against the CAZy with DIAMOND blastx (e-value threshold of 1×10−5).

### Bordetella pertussis strains annotation

We annotated the B. pertussis strains involved in the whooping cough outbreaks since 2000 using the outbreak time information collected from previous studies^[31, 32]^. Based on the results of Ring et al., we defined two major outbreak periods: before and after 2011.

### The DeepInverton framework

#### Curating training and testing data

In core dataset, the sequences identified to be invertons by PhaseFinder were determined as the inverton data, and the rest was categorized to the non-inverton data. We analyzed the density distribution of the length of the 10,983 inverton sequences in the core dataset, and found that more than 75% of the sequences were shorter than 300 bases. So we finally selected the inverton sequences shorten than 300 bases as the positive inverton data. To attenuate the impact of uneven distribution of the number of species on the training of deep learning models, a similar number of non-inverton sequences were randomly obtained as the negative inverton data. Training and 5-fold cross-validation of DeepInverton-core were performed using 75% of the total dataset. The remaining 25% of samples were reserved as an external test set to evaluate the model performance. The train-test split was stratified by label to ensure that each dataset maintained a label distribution representative of the entire dataset (50% positive inverton sequences and 50% negative inverton sequences). For DeepInverton-extended, the training dataset included 133,617 inverton sequences and a similar number of non-inveron sequences obtained from the extended dataset. The model performance was also evaluated using the external test set.

#### Deep Learning Model Architecture and Hyperparameter Tuning

Deep learning models were implemented by Python 3.9.13 using standard libraries that are publicly available: Pytorch (1.13.0) and Scikit-learn (1.1.2). Hyperparameter tuning was used to identify the best model architectures and parameters. For the final DeepInverton model, there were 6 layers: an input layer, an one-hot encoded vector with dimensions of 300 × 5, three hidden layers with the RELU activation function, and an output layer with a softmax function. A cross-validation procedure was applied to determine the within-training set performance by splitting data into training and validation sets for fivefold-stratified cross-validation. The optimal models selected based on cross-validated results were then evaluated in the external test dataset and AUROC value was calculated accordingly for the visualization of results.

### Models comparison

Random forest (RF) models were implemented with the following parameters: n_estimators=500 and max_leaf_nodes=16. For BLAST, it was performed using BLASTn (v2.13.0) with the default parameters. The positive inverton sequences in the training set were selected to build the target database, and the sequences in the test set which were successfully aligned to the database were identified as the predicted inverton sequences. In order to ensure data consistency, we adopted the same training and test set as the DeepInverton. For model evaluation, we included AUROC and AUPR to characterize the model performance.

#### Metagenome mining with DeepInverton

We initially used the einverted tool from EMBOSS^[49]^ to search for sequences flanked by reverted repeats in the metagenome data, filtering out sequences with a length less than 300 base pairs. Subsequently, we fed these sequences into the DeepInverton model and obtained the probability of the particular sequence to be an inverton or not. Based on the performance of the DeepInverton on external test set, we observed a significantly low false positive rate when the logarithm (base 10) of the ratio between the positive inverton probability and the negative inverton probability reached 15. To ensure high fidelity in the identification of inverton sequences in the metagenomes, we considered sequences as final inverton candidates only if their logarithm (base 10) ratio of positive inverton probability to negative inverton probability exceeded 15.

#### Antibiotic resistance experiments

The 23 candidate invertons and 3 non-invertons identified by DeepInverton were synthesized with their downstream antibiotic resistance genes by GENEWIZ (Suzhou, China). Both ON and OFF orientation inverton sequences were synthesized to represent the regulatory forms of the invertons in vivo. Also, ARGs with known promoters were synthesized as controls for antibiotic resistance experiments. All the produced sequences were verified by sanger sequencing and introduced into E. coli. These strains were then streaked on Luria-Bertani (LB) agar medium and incubated at 37 °C overnight. The individual colonies were picked into LB culture medium and shaken at 220 r.p.m. at 37 °C overnight. The LB bacterial suspension was diluted to the predetermined starting concentration (optical density at 600nm (OD600=0.1)) and then again diluted 1,000 times for the antibiotic resistance test. We dissolved the antibiotic powder in ultrapure water or dimethyl sulfoxide (DMSO) according to its solubility and diluted it to 1 μg/mL. We set four experimental groups to test the candidate invertons’ (including the 3 candidate non-invertons’) regulatory activity: (1) positive control group, 4 mL bacterial solution (with known promoter and ARG) and 80 μl antibiotic solution; (2) negative control group, 4 mL bacterial solution (without ARGs) and 80 μl antibiotic solution; (3) inverton-ON group, 4 mL bacterial solution (with ON-orientation inverton and ARG) and 80 μl antibiotic solution; (4) inverton-ON group, 4 mL bacterial solution (with OFF-orientation inverton and ARG) and 80 μl antibiotic solution. The OD600 value of each group was measured after culture at 37 °C for 12 h. We used R software to perform statistical test for comparing experimental groups with the control groups. All experiments were performed with three independent replicates.

#### MIC determination

Minimal inhibitory concentrations (MICs) of the antibiotic for the strains containing resistance genes were determined using E-tests. In brief, E.coli strains were inoculated in Luria-Bertani (LB) medium at 37 °C overnight. The cultures were diluted 1:100 using the fresh LB and subsequently cultured to the exponential phase (OD600 of 0.4-0.6), and then the cell concentration was diluted to an OD600 of 0.1, and 200 μl bacterial solution were spread evenly onto LB agar medium. The corresponding antibiotic MIC test strips were then sticked to the center of the medium. After incubating at 37 °C for 16-18h, the MIC was determined as the minimum concentration of antibiotic where bacteria showed no detectable growth. All experiments were performed with three independent replicates.

## Data availability

The Inverton database of more than 200,000 candidate invertons with detailed genomic information is available at http://inverton-db.aimicrobiome.cn/. The DeepInverton model has also been made publicly accessible at https://github.com/HUST-NingKang-Lab/DeepInverton. Source data of invertons (xxx) is available for download via Zenodo.

## Acknowledgements

We thank Hong Bai and Experimentation teaching center of the college of life science and technology for their assistance with antibiotic resistance experiments. Numerical computations were performed on the Hefei Advanced Computing Center.

